# Reconstructing species evolutionary trees using the numerical features of microRNA

**DOI:** 10.1101/770503

**Authors:** Rongsheng Zhu, Dawei Xin, Zhanguo Zhang, Zhenbang Hu, Yang Li, Qingshan Chen

## Abstract

Research has revealed that some microRNAs show obvious lineage or species specificity, but others show highly conserved properties among species. Based on these properties, we aimed to reconstruct a species evolution tree using a new technique that refers to the numerical features of microRNA. First, we selected 132 microRNA numerical features that included base content, secondary structure matching state frequency, free energy features and information entropy features, and 32 species that included 22 animals, 9 plants and a representative virus group. Second, we found several significantly different numerical features among lineages or species by statistical analysis and confirmed that differences in each numerical feature were not identical. Third, we designed a comprehensive feature and confirmed that it showed obvious lineage and species specificity. Last, species trees were built using the comprehensive feature. The results showed that the reconstructed species tree was almost in keeping with the actual chronological order of species evolution. This indicated that our analysis was effective. Our research strategy offers a new route for investigating species evolution.

## Introduction

MicroRNAs are a class of non-coding small RNAs that act as important regulators after transcription. Their length is about 19–22 nt and they function by degrading or suppressing target genes [1-4]. In recent years, much research [5-13] has been carried out on their functions and the results have made the functions of many kinds of microRNA increasingly clear. However, there has been much dispute about the evolution and origin of microRNA [14]. A lot of research on microRNA’s evolution and origin has been conducted by investigating species evolution [15-19] and family evolution [20-22]. The results showed that there is species or lineage specificity in microRNA sequences, secondary structure and distribution among animals or plants. Certainly, there are many specific and non-specific microRNA genes and this specificity is just a result of evolutionary selection [14]. This conclusion supplied a theory base for investigating species evolution through all of the microRNA genes of a certain species. However, there were no tools or methods available to test our idea. Fortunately, the results of one study [23] showed that microRNA numerical features possessed species-specificity, which we used as a tools base for investigating species evolution through all of the microRNAs of a species. We selected 32 animals and plants as research materials and extracted 132 numerical features that included sequence, structure, energy and information entropy. First, we examined the ability of numerical features to distinguish species. Second, we evaluated species relationships by a directed statistic feature. Third, we designed a comprehensive numerical feature and confirmed that it possessed significant species-specificity. Last, we reconstructed animal and plant evolutionary trees based on the comprehensive numerical feature and the results showed that the reconstructed animal tree was almost consistent with real species evolution time. Thus, we confirmed that our idea was feasible and our analysis strategy reasonable. Our research strategy offers a new route for investigating species evolution.

## Materials and Methods

### Choice of species

Thirty-two species that had over 100 microRNAs in mirBASE (Version 17) [37] were selected, which covered animals, plants, and viruses and included 10886 microRNA genes. All microRNAs from viruses were considered as the same class. In addition, two classes were chosen to verify the research results, one was primates, which included *Homo sapiens, Macaca mulatta, Pan troglodytes*, and *Pongo pygmaeus*, and the other was dicotyledonous plants, which included *Arabidopsis thaliana, Glycine max, Medicago truncatula, Populus trichocarpa*, and *Vitis vinifera*. Basic information for the microRNAs of the candidate species is shown in Table 1.

### Feature selection

One hundred and thirty-two numerical features in eight classes were selected in total; the serial numbers and names of the features are described in Supplementary Table 1. The first class referred to the frequency characteristics of single nucleotides. The second class referred to two-base combinations of the four bases, A, C, G, and U. The third class referred to three base combinations of the four bases.

The fourth class referred to frequency features of the secondary structure matching state. Based on RNA secondary structure predicted by Mfold [38], the matching state of each nucleotide was described using the method presented by Xue et al. [39]. For example, “A+.” indicates that the nucleotide at the site is “A”, with a non–matching left site and a non-matching right site in the secondary structure. Examples are shown in Fig. 1. In total, there were 32 frequency features for the secondary structure matching state.

**Fig. 1.**
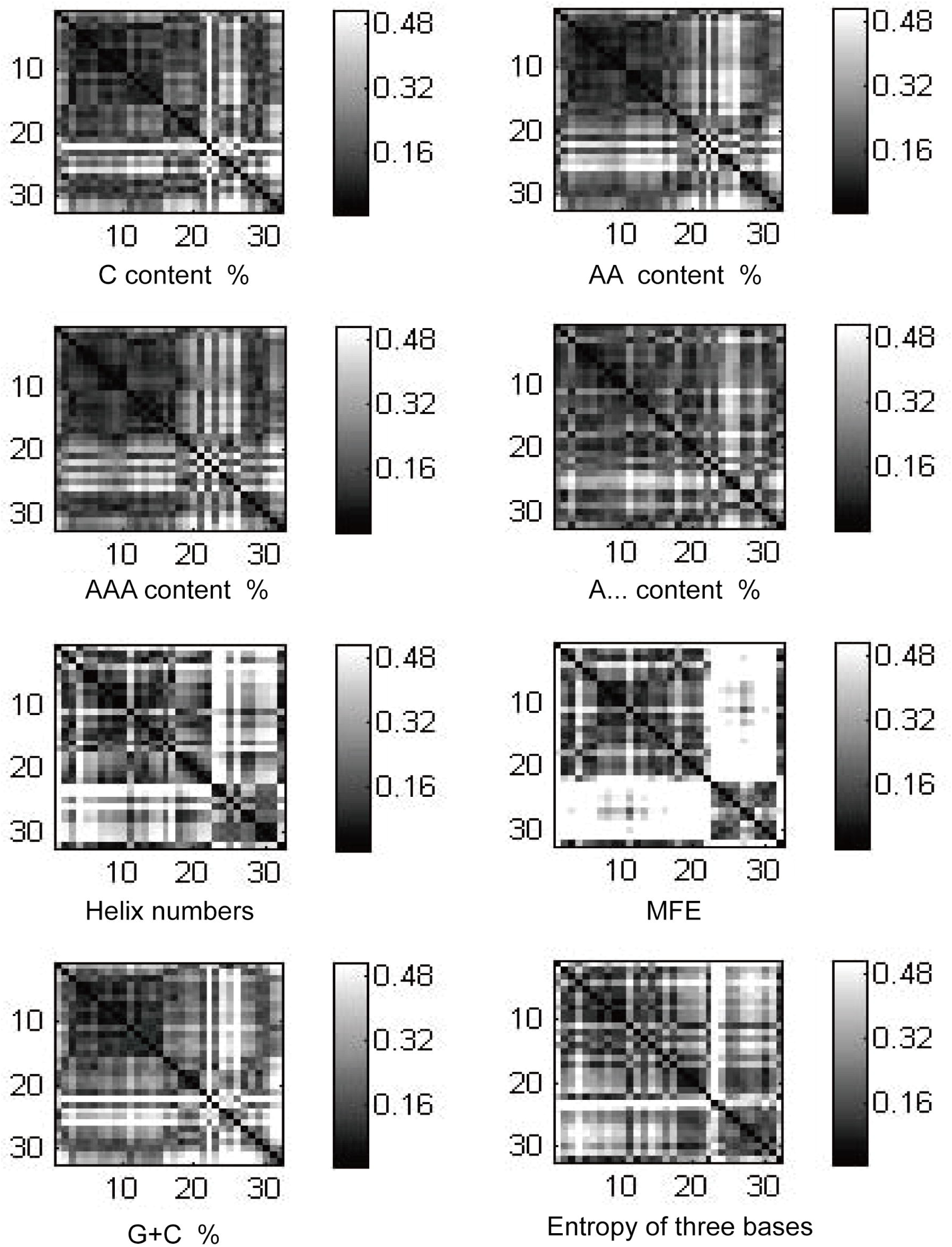
Eight grayscale maps based on eight numerical features among 32 species. The eight subplots showing the distribution of differences for eight features of the 32 species were drawn based on Kolmogorov-Smirnov test statistics. From top to bottom and from left to right, the numbers in the figure refer to the candidate species: 1 - *Ciona intestinalis*, 2 - *Xenopus tropicalis*, 3 - *Gallus gallus*, 4 - *Canis familiaris*, 5 - *Equus caballus*, 6 - *Monodelphis domestica*, 7 - *Macaca mulatta*, 8 - *Homo sapiens*, 9 - *Pan troglodytes*, 10 - *Pongo pygmaeus*, 11 - *Ornithorhynchus anatinus*, 12 - *Mus musculus*, 13 - *Rattus norvegicus*, 14 - *Bos taurus*, 15 - *Sus scrofa*, 16 - *Danio rerio*, 17 - *Fugu rubripes*, 18 - *Bombyx mori*, 19 - *Drosophila pseudoobscura*, 20 - *Caenorhabditis elegans*, 21 - *Capitella teleta*, 22 - *Schmidtea mediterranea*, 23 - *Physcomitrella patens*, 24 - *Arabidopsis thaliana*, 25 - *Glycine max*, 26 - *Medicago truncatula*, 27- *Populus trichocarpa*, 28 - *Vitis vinifera*, 29 - *Oryza sativa*, 30 - *Sorghum bicolor*, 31 - *Zea mays*, 32 - Virus. The eight features represent the eight feature regions that are detailed in Supplementary Table 1.

The fifth class included the length of microRNA genes, the number of bulge loops, the number of helices, the number of interior loops, and the number of stacks. Except for the length of the gene, the features were taken from Mfold predictions of secondary structure. Detailed examples are shown in Fig. 1. The sixth class included the minimum free energy (MFE) [40], the adjusted minimum free energy (AMFE) [41], and the minimum free energy index MFEI [42]. The seventh included G+C content, (G+C)/(A+U) ratio, A/C ratio, and G/U ratio.

The eighth class referred to features with a relationship to information entropy. The information entropy [43] was calculated using the formula:

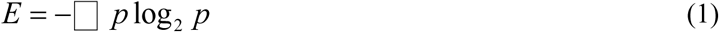

Formula (1) generated four kinds of entropy information related to the frequency of single nucleotides, dual nucleotides, triple nucleotides, and the matching state frequency of the secondary structure (termed Sec_str_entr in Supplementary Table 1).

The first to eighth classes were designated A, B, C, D, E, F, G and H in corresponding order.

### Methods

#### Feature Identification Number of species pairs (FIN)

The Feature Identification Number of species pairs was the number of candidate features that could be used to distinguish a certain species pair, i.e. there was a significant difference between the species pair for a certain feature according the Kolmogorov-Smirnov test. The formula for Feature Identification Ratio (FIR) of species pairs was defined as follows:

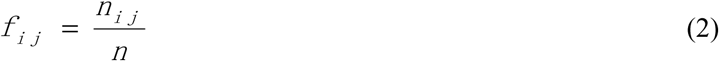

In formula (2), *n* denotes the total number of candidate features, and *n*_(*i,j*)_denotes the feature identification number between species *i* and species *j*. In this study, FIN and FIR were used to measure the degree of genetic relationship between species.

FIN could effectively reveal the genetic relationships of species. However, FIN and FIR showed reverse changes in the extent of genetic relationships of species. Thus, Similarity Identification Degree (SID) was used as a consistent measure of the genetic relationships of species. SID was defined as follows:

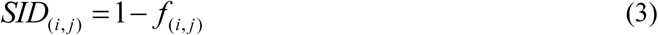

In formula (3), *i* and *j* denote sequence numbers of species and *f*_(*i,j*)_ denotes the FIR of species *i* and species *j*. From the definition of SID, the bigger the SID value, the closer the genetic relationship of the species.

#### Species Pairs Identification Number of a feature (SPIN)

The Species Pairs Identification Number refers to the number of species pairs that are distinguished by a certain feature. Subsequently, the Species Pairs Identification Ratio of a feature (SPIR) was defined as follows:

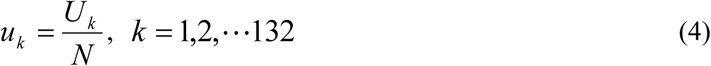

In formula (4), *k* denotes the sequence number of the feature, *U*_*k*_ denotes the SPIN value for feature *k*, and *N* denotes the total number of species pairs. SPIR could describe the ability of a feature to identify a species in a background of species pairs and was used to evaluate candidate features.

#### Feature Difference Map (FDM)

FIR and SID were both able to comprehensively evaluate genetic relationships for a pair of species based on their background features information. However, they involved counting numbers of significantly different features between species, and could not display the extent of difference for every feature between species. To assess the detailed differences and similarities between species for a single feature, a Feature Difference Map (FDM) was designed based on certain features of the microRNAs. The FDM is a grayscale map and the intensity of each spot is determined using the Kolmogorov-Smirnov test statistic for the feature between two species. Kolmogorov-Smirnov’s test statistic [33] is:

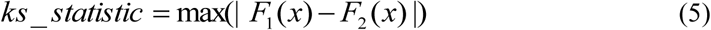

In formula (2), *F*_1_(*x*) is the empirical distribution function of the first sample and *F*_2_(*x*) is the empirical distribution function of the second sample.

#### Integrated Numerical Feature (INF)

We analyzed the data sets, which included 10886 microRNA genes and 132 numerical features, and obtained 10886 row and 132 column PCA scores by principal component analysis. The results were recorded as *PCA_Score* = (*x*_*ij*_),(1,…10886; *j* = 1,…132). Meanwhile, 132 eigenvalues were obtained by PCA and were marked as *λ*_*i*_(*i* − 1,…132).

The Integrated Numerical Feature (INF) was defined as follows:

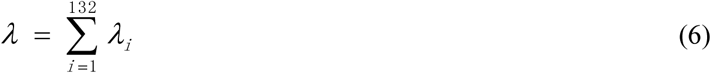

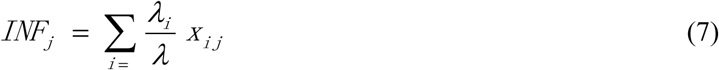

The INF was a proper feature that could effectively eliminate linear dependence among numerical features.

## Results

### Differences and similarities among species based on each numerical feature of microRNA (Fig. 1)

To discover numerical feature changes based on different species, eight features (representing classes A–H) were selected from the 132 candidate numerical features of microRNAs to create FDMs. The eight features were the C base frequency, AA base pairs frequency, AAA base frequency, A… frequency, helix numbers, MFE, G+C content and trinucleotide entropy.

In Fig. 1, the classifications in the eight FDMs were roughly consistent with the biological classes, but some were clearer, including helix numbers, MFE, G+C content and trinucleotide entropy, while others were vague. This illustrated that numerical features of microRNA were related to species classification and that the extent of the relationship differed. One principal reason for the differing extents of the relationships could be that different features represent different biological properties. From several of the clearer FDMs, we found that microRNA genes possessed high species specificity that could be reflected by numerical features. These FDMs offered a way to investigate species evolution through the numerical features of these small molecules.

To reveal the ability of each numerical feature to distinguish species pairs, Fig. 2 was created based on SPIR values. The eight classes of features (A–H) are shown from left to right in Fig. 2. For the four single nucleotides (class A), the identification ratio among the features was consistent, and the average identification ratio was 60.13%. For the sixteen dual nucleotide features (class B), the identification ratio among the features showed significant differences, with an average identification ratio of 47.99%. For the 64 triple nucleotide features (class C), the identification ratios among the features were moderately consistent, and the average identification ratio was 45.01%. For the thirty-two secondary structure pairing state features (class D), the feature identification ratios were lower than for the other classes, with an average identification ratio of 30.31%. This result demonstrated that there were no significant differences among the secondary structures of the microRNAs from all species. For the length and direct counting numbers of secondary structures (class E), the feature identification ratio of length reached 89.72%, which was the highest of all the features. The feature identification ratios for the number of stacks, number of helices, number of interior loops, and number of bulge loops were 73.39%, 65.12%, 51.81%, and 14.31%, respectively. These results showed that there was generally a difference in the length of microRNA genes among species, but there was little difference in the number of bulge loops. For the features related to minimum free energy (class F), the feature identification ratios for MFE, AMFE, and MFEI were 75.6%, 61.9%, and 60.69%, respectively, which showed relatively high identification ratios, and a bigger change in features among species. For base content and base content ratios (class G), a high average identification ratio of 68.85% was observed. For the four entropy-related features (class H), the feature identification ratio of the entropy of triple-nucleotides was 70.36%, which was the highest in class H, and was less than 50% for the other features.

**Fig. 2.**
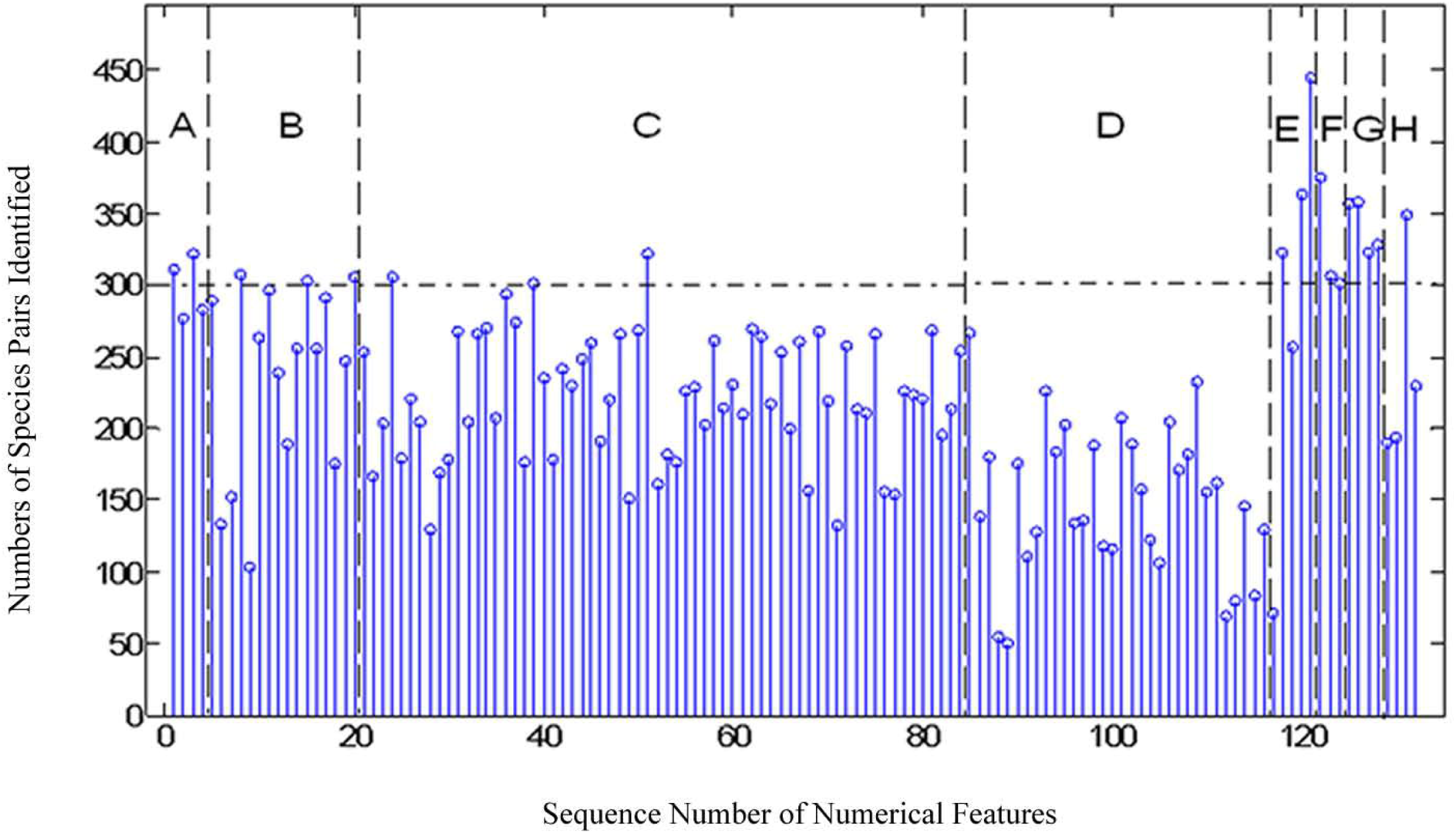
SPIN values of 132 numerical features of microRNAs. Species Pairs Identified Numbers were calculated for 496 pairs of species based on 132 numerical features. The figure is divided into eight regions (A–H). These regions are described in Supplementary Table 1.

Overall, the top six for recognition rate among the selected 132 features were the sequence length (Length, 89.72% recognition rate), the minimum free energy (MFE, 75.60%), the number of stacks (Stack, 73.39%), (G + C)/(A + U) (72.18%), G + C (71.96%), and the entropy of triple nucleotide content (70.36%). There were significant differences among species identified with these features and they tended to be specific for species. In contrast, the bottom six features for recognition rate were A.++ (10.08%), A+.. (11.08%), U+. (13.91%), Bulge loop number (14.31%), U.++ (16.13%), and U++. (16.94). There was little difference among species identified with these features and they tended to be properties of individual microRNAs themselves and not lineage- or species-specific.

### Differences and similarities among species based on all numerical features of microRNA (Fig. 3)

**Fig. 3.**
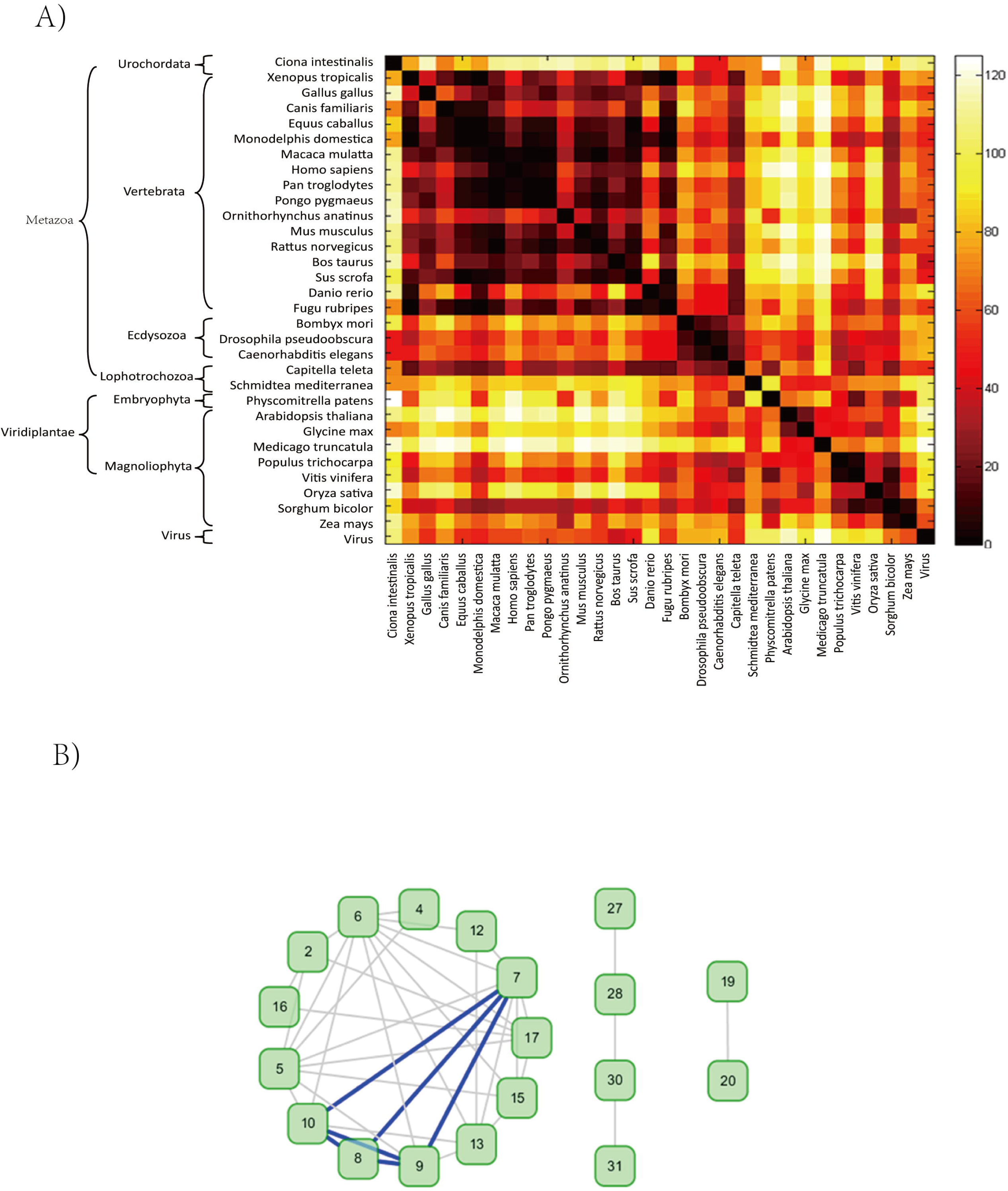
Species relationships based on Feature Identification Numbers (FINs). (A) A hot map that demonstrates the state of 32 species identified by the FINs for 132 features. From the bottom to the top of the color bar in the right of graph, the color changes from dark to light represent FIN changes from 0 to 128. The bigger the FIN, the lighter the grid square and the more distant the relationship between the two species. (B) A relationship graph based on species pairs difference feature numbers less than 9. The numbers in the figure represent the following candidate species: 2 - *Xenopus tropicalis*, 4 - *Canis familiaris*, 5 - *Equus caballus*, 6 - *Monodelphis domestica*, 7 - *Macaca mulatta*, 8 - *Homo sapiens*, 9 - *Pan troglodytes*, 10 - *Pongo pygmaeus*, 12 - *Mus musculus*, 13 - *Rattus norvegicus*, 15 - *Sus scrofa*, 16 - *Danio rerio*, 19 - *Drosophila pseudoobscura*, 20 - *Caenorhabditis elegans*, 27 - *Populus trichocarpa*, 28 - *Vitis vinifera*, 30 - *Sorghum bicolor*, 31 - *Zea mays*. There appear to be three clusters in the graph. The first cluster includes 19 and 20, which belong to Ecdysozoa. The second cluster includes 27, 28, 30, and 31, which belong to Magnoliophyta. The third cluster includes the others, which belong to Vertebrata. Species 7, 8, 9, and 10 in the third cluster belong to the Primates and the species pairs difference feature numbers within them are less than 4.

To survey how all 132 numerical features performed in distinguishing species as a whole body, Fig. 3 was created based on FIR and SID values.

#### Species difference map

A 32 × 32 matrix was constructed for the 32 species from the species classification order. The definitive element of the matrix was the SPIN between two corresponding species. The matrix was named the species difference matrix (SDM). Furthermore, based on the SDM, a hot map was drawn as a species difference map (SDMP) (Fig. 3A).

In Fig. 3A, the dark square regions in the upper-left and lower-right corners indicated that there were few differences in microRNA features among animals or plants. Moreover, the upper-right and lower-left corners of the map showed brighter colors, which indicated that there was a clear difference in terms of microRNA features between animals and plants. The greater the genetic distance among species, the brighter the color of the elements in the matrix, which provided strong evidence that FIN values were able to describe the genetic relationships of species pairs.

There were very few differences between *Ciona intestinalis* from Urochordata and *Bombyx mori, Drosophila pseudoobscura*, and *Caenorhabditis elegans* from Lophotrochozoa; however, there were more differences between *Ciona intestinalis* and the other species, which indicated that there was a closer genetic relationship between Urochordata and Lophotrochozoa than either of them had with the other species. Similarly, there were clear dark squares among the three animal species in *Lophotrochozoa*, indicating a high similarity among species in the same class, but there were fewer SPINs between the species of the Lophotrochozoa and *Xenopus tropicalis, Danio rerio*, or *Fugu rubripes* from Vertebrata, and *Capitella teleta* from Lophotrochozoa.

In the sixteen species from Vertebrata, there was apparently a difference between *Danio rerio* and the other species, while the FIN values among the other species, except *Danio rerio*, were very small, which showed that microRNA genes are highly conserved among Vertebrata species and among animal species as a whole. Furthermore, a smaller FIN value was obtained between *Danio rerio* and *Fugu rubripes* than with other species, which showed that there was high conservation within species.

For plants, the matrix was darker among the species in Magnoliophyta, but brighter between Magnoliophyta and the other classes. The strength of the color was apparently inferior to that of Vertebrata. These results showed that there were bigger differences among the microRNA genes of species in Magnoliophyta, which supported the hypothesis that there is great diversity in the microRNA genes of plants. The matrix also showed that viral microRNA genes were more similar to those of animals than plants.

In summary, the SDMP constructed using FINs could directly display the degree of difference between species pairs. The SDMP could display basic differences and similarities between species or classes, or within classes. Thus, SDMP could be used as a valid tool to display the genetic relationships of species.

#### Species relationship map

Because SDMP or FIN was a reverse description, SID was introduced. A species relationship map (Fig. 3B) was created based on SID.

The threshold for SID was set as 0.8. The candidate species were divided into four classes: animals, plants, viruses, and *Urochordata*. In Fig. 3B, *Bombyx mori, Drosophila pseudoobscura, Caenorhabditis elegans, Capitella teleta*, and *Schmidtea mediterranea*, which have similar shapes or habits, were classified into the same group by SID analysis. All primates were classified into the same group as *Homo sapiens. Glycine max, Arabidopsis thaliana*, and *Medicago truncatula*, belonging to Magnoliopsida, were classified into the same sub-class; *Oryza sativa, Sorghum bicolor*, and *Zea mays*, belonging to Liliopsida, were classified into another sub-class. These results showed that SID could reflect the genetic relationships among species.

### Differences and similarities among species based on the Integrated Numerical Feature of microRNA (Fig. 4)

**Fig. 4.**
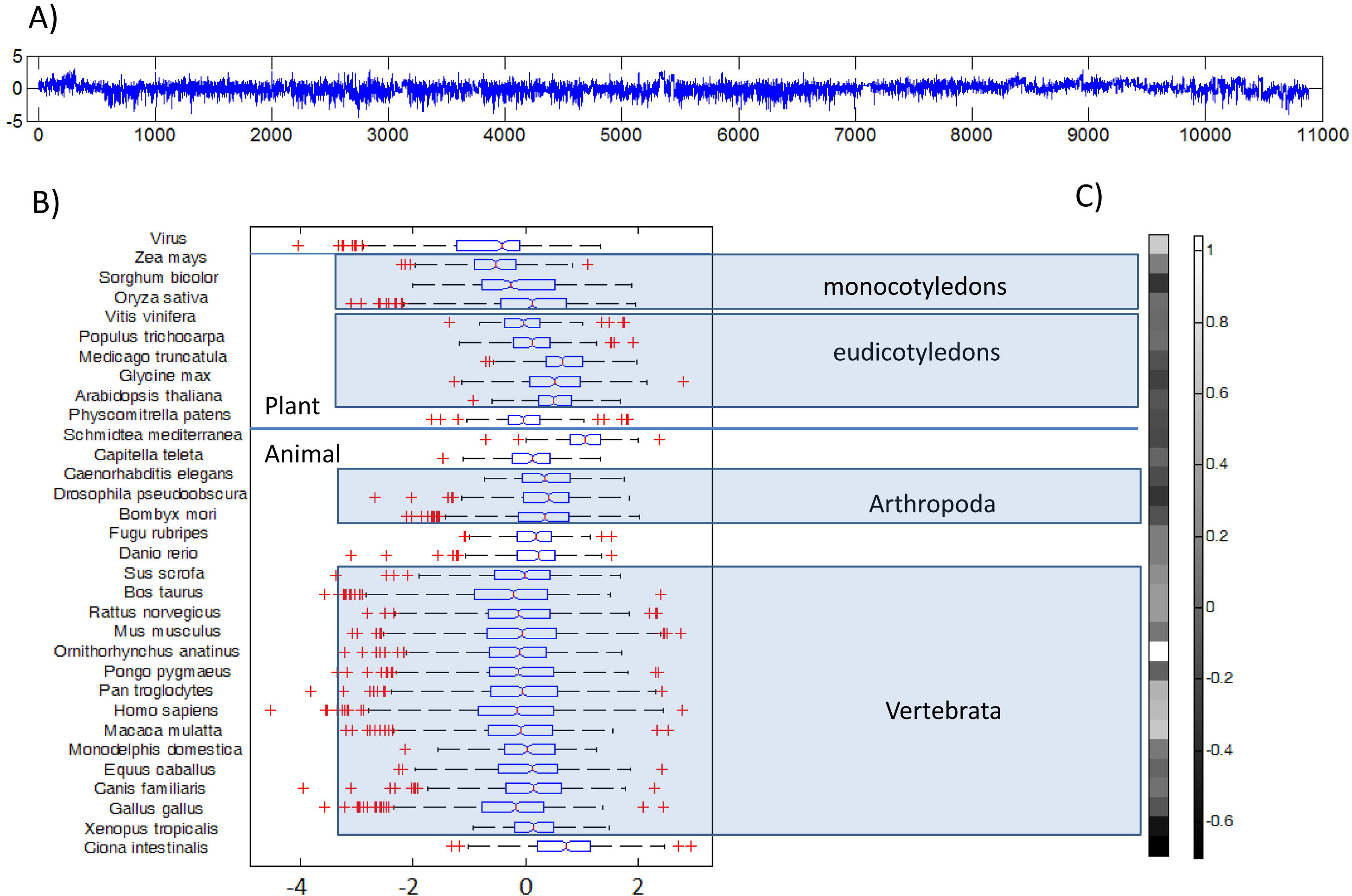
Distribution of the microRNA comprehensive numerical feature. (A) The basic distribution of 10886 microRNA’s INF values. (B) Boxplots for every species based on INF values. (C) A grayscale map that was drawn based on the mean INF values for each species.

Although FIN and SID might describe the relationship between species, linear dependence among features made the relationship imperfect. To overcome this, INF was introduced. Fig. 4 was drawn based on INF, which was an integrated numerical feature of microRNA. Fig. 4A shows that there were more significant differences among species based on INF than FIN. This illustrated that the numerical features of microRNAs possessed obvious species specificity.

To clarify the overall level of INF based on different species, Fig. 4B and 4C were created. In Fig. 4B, there were significant differences between animals and plants based on INF and these differences were still obvious within the plant or animal groups. In animals, there was a significant difference between Arthropoda and Vertebrata, but the INF values were remarkably consistent in the interior of each. In plants, although there were differences between Monocotyledons and Eudicotyledons for INF, there were still differences within the interior of each group. These results indicated that the microRNA genes of animals possessed more highly conserved classification properties than those of plants, and that the microRNA genes of plants had various changes based on species class. In Fig. 4B, the virus group and *Ciona intestinalis* were of particular interest. The INF of the virus group tended to be at minimum and the INF of *Ciona intestinalis* tended to be at maximum.

Fig. 4C is a grayscale map that was generated based on the mean INF values of each candidate species. There were significant differences among different species in this figure. From these three figures, we realized that INF might become a valuable tool to measure the relationships between species and the result made us interested to build a species relationship map to discover whether the numerical features were closely related to the classification and evolution of species.

### Building species evolutionary relationships based on the Integrated Numerical Feature (Fig. 5)

**Fig. 5.**
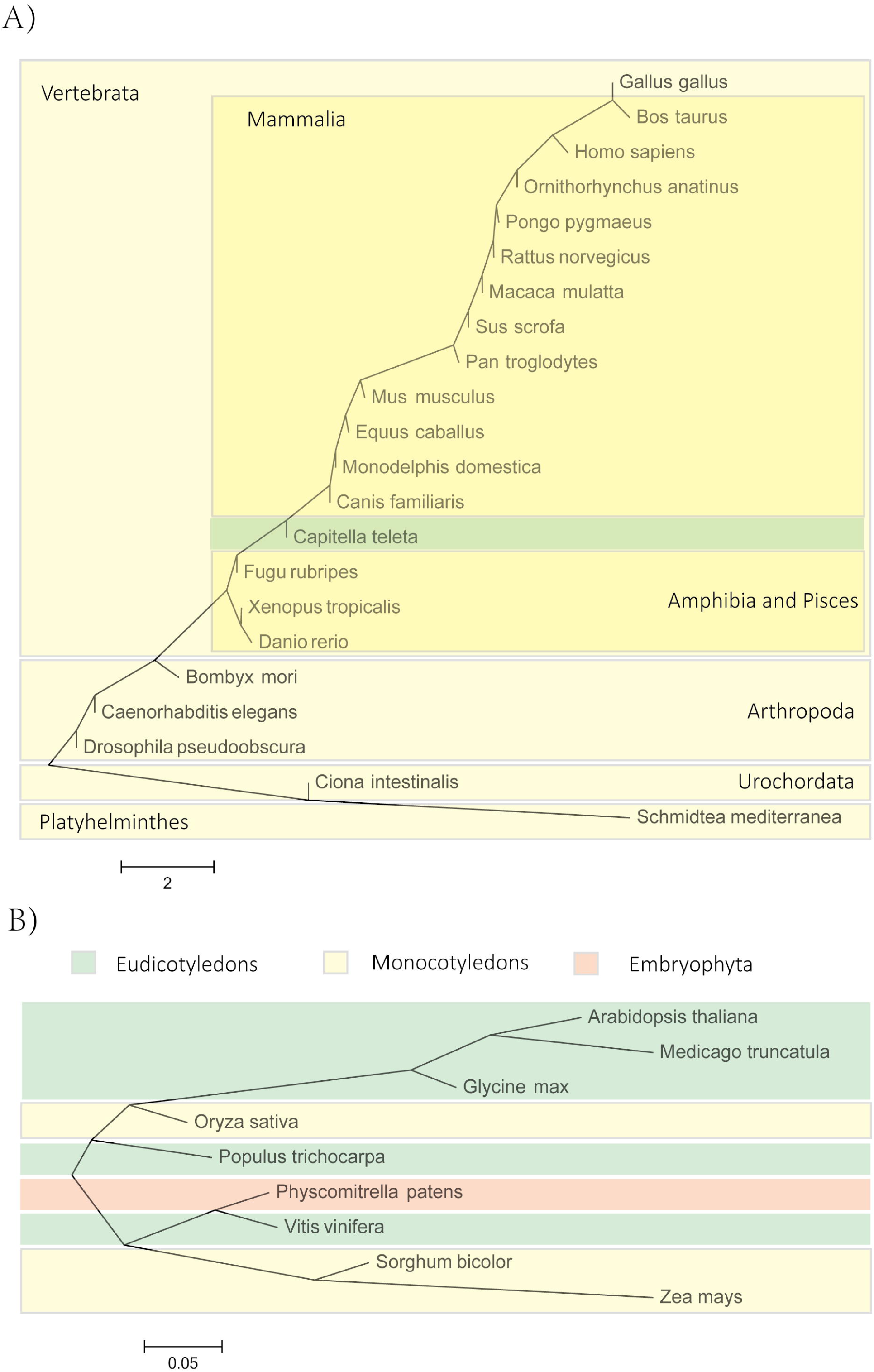
Species trees reconstructed for animals and plants based on INF values. (A) Animal species evolution tree. (B) Plant species evolution tree.

Two species relationship trees were built. One was for animals, the other for plants. Measurement of each two-species relationship was by a *t*-test statistic between the two species’ INF values. The trees were drawn using the MEGA [24] software.

In Fig. 5A, the candidate animal species were classified and the results based on INF were almost in accordance with the actual classification, except for *Capitella teleta.* This demonstrated that animal microRNA genes possessed obvious species class aggregation, which had been confirmed by the results of the latest research [14]. This type of tree may become an option for investigating animal evolution and species classification at the molecular level.

On the contrary, in Fig. 5B, the evolutionary tree that was built based on INF was not in accordance with the actual evolution order or species classification. The reason for this may be that plant microRNA genes possess more complex structures and more variable nucleotide sequences than those of animals [14].

## Discussion

As more and more microRNAs are found, many studies on them are being carried out, for example on their functions, origins, identification and evolution. Much research has confirmed that some microRNAs are highly conserved, while other novel miRNAs are very much actively evolving. Obviously, the highly conserved microRNAs are related to some core life processes and most of the novel ones are related to species-specific functions. In some reports, some of the numerical features of microRNAs were related to species classes, which showed that the numerical features could indicate microRNA conservation and specificity. In our analysis, we investigated species evolution and classification by a new route using the numerical features of microRNAs.

In some studies [25-31], relationships were found between the numerical features of microRNAs and species class or evolution. We have proved these relationships based on an analysis of 32 species by classical statistics methods, novel measures and microRNA numerical features. In concrete terms, most features displayed significant differences between species classes. However, some of them were weaker than others, and some were stronger, which made us unable to use FIN or FIR as a measure for evaluating the weakness or strength of relationships between species. To obtain a reasonable measure to evaluate the relationships of species, we designed a comprehensive feature – the Integrated Numerical Feature (INF). The INF effectively compensated for the deficiencies of FIN and FIR, which derived from accumulating single numerical feature data to evaluate differences in species. Finally, we used INF to rebuild species phylogenetic trees.

Of the species trees we built, the animal tree was closer to the real species classification than the plant tree. The reason for this may be that the pre-microRNA of animals was more conserved than that of plants, and the sequence and secondary structure of plant microRNAs have become more varied than those of animals [14]. In the animal tree, we found that the species order on the tree was very close to the fossil record time [32, 33]. Deuterostomia, which includes *Bombyx mori, Caenorhabditis elegans* and *Drosophila pseudoobscura*, originated about 555 MYA (million years ago) [34]. *Ciona intestinalis* of Urochordata originated between 548 and 555 MYA [35]. Amphibia and Pisces originated between 485 and 541 MYA and *Capitella teleta* belonging to the annelids originated about 505 MYA. Mammalia originated around 167 or 225 MYA; there has been debate about these two time points [36]. Our results showed that the microRNA genes of animals possessed obvious species or lineage specificity, that INF had the ability to distinguish species classes, and that our research strategy might be used to build species trees using other DNA or RNA molecular classes to discover different perspectives on species evolution. Although the high consistency between the rebuilt tree and fossil times in animals was not observed in plants, three eudicotyledonous species that included *Arabidopsis thaliana, Medicago truncatula* and *Glycine max* and two monocotyledonous species that included *Sorghum bicolor* and *Zea mays* kept their species classification aggregation. The plant evolution order and classification predicted for the others were not ideal. The reason for this might be the amount of variability among plant species in their microRNA genes or it might be related to the selection of microRNA numerical features. In any case, we confirmed that our supposition was feasible. This offers a new route for investigating species evolution using the numerical features of many kinds of molecules.

## Abbreviations

MFE: minimal free energy;
AMFE: adjusted minimal free energy;
MFEI: minimal free energy index;
FIN: feature identification number;
FIR: feature identification ratio;
SID: similarity identification degree;
SDM: species difference matrix;
SDMP: species difference map;
SID: similarity degree of identification;
SPIN: species pairs identification number of feature;
SPIR: species pairs identification ratio of feature;
FDM: features difference map;
INF: integrated numerical feature.

## Tables

**Table 1.** Basic information on candidate species’ microRNAs.

## Supporting information

**Supplementary Table 1.** Performance of 132 numerical features in identification of species pairs.

**Supplementary Table 2.** Statistics for species pairs being investigated.

