## Supplementary Table 1 for "Reconstructing species evolutionary trees using the numerical features of microRNA"

Table S1 Performance of 132 numerical features in identification of species pairs

| Region | Sequence Number | Content | FIN | FIR |
| --- | --- | --- | --- | --- |
| A | 1 | A | 311 | 0.627016129032258 |
|  | 2 | C | 277 | 0.558467741935484 |
|  | 3 | G | 322 | 0.649193548387097 |
|  | 4 | U | 283 | 0.570564516129032 |
| B | 5 | AA | 290 | 0.584677419354839 |
|  | 6 | AC | 133 | 0.268145161290323 |
|  | 7 | AG | 152 | 0.306451612903226 |
|  | 8 | AU | 308 | 0.620967741935484 |
|  | 9 | CA | 103 | 0.207661290322581 |
|  | 10 | CC | 263 | 0.530241935483871 |
|  | 11 | CG | 297 | 0.598790322580645 |
|  | 12 | CU | 239 | 0.481854838709677 |
|  | 13 | GA | 189 | 0.381048387096774 |
|  | 14 | GC | 256 | 0.516129032258065 |
|  | 15 | GG | 303 | 0.610887096774194 |
|  | 16 | GU | 256 | 0.516129032258065 |
|  | 17 | UA | 291 | 0.586693548387097 |
|  | 18 | UC | 176 | 0.354838709677419 |
|  | 19 | UG | 247 | 0.497983870967742 |
|  | 20 | UU | 306 | 0.616935483870968 |
| C | 21 | AAA | 253 | 0.510080645161290 |
|  | 22 | AAG | 167 | 0.336693548387097 |
|  | 23 | AAC | 204 | 0.411290322580645 |
|  | 24 | AAU | 306 | 0.616935483870968 |
|  | 25 | ACA | 179 | 0.360887096774194 |
|  | 26 | ACG | 221 | 0.445564516129032 |
|  | 27 | ACC | 205 | 0.413306451612903 |
|  | 28 | ACU | 130 | 0.262096774193548 |
|  | 29 | AGA | 169 | 0.340725806451613 |
|  | 30 | AGG | 178 | 0.358870967741936 |
|  | 31 | AGC | 268 | 0.540322580645161 |
|  | 32 | AGU | 205 | 0.413306451612903 |
|  | 33 | AUA | 266 | 0.536290322580645 |
|  | 34 | AUG | 271 | 0.546370967741936 |
|  | 35 | AUC | 207 | 0.417338709677419 |
|  | 36 | AUU | 294 | 0.592741935483871 |
|  | 37 | GAA | 274 | 0.552419354838710 |
|  | 38 | GAG | 177 | 0.356854838709677 |
|  | 39 | GAC | 301 | 0.606854838709677 |
|  | 40 | GAU | 235 | 0.473790322580645 |
|  | 41 | GCA | 178 | 0.358870967741936 |
|  | 42 | GCG | 242 | 0.487903225806452 |
|  | 43 | GCC | 230 | 0.463709677419355 |
|  | 44 | GCU | 249 | 0.502016129032258 |
|  | 45 | GGA | 260 | 0.524193548387097 |
|  | 46 | GGG | 191 | 0.385080645161290 |
|  | 47 | GGC | 220 | 0.443548387096774 |
|  | 48 | GGU | 266 | 0.536290322580645 |
|  | 49 | GUA | 151 | 0.304435483870968 |
|  | 50 | GUG | 269 | 0.542338709677419 |
|  | 51 | GUC | 322 | 0.649193548387097 |
|  | 52 | GUU | 161 | 0.324596774193548 |
|  | 53 | CAA | 182 | 0.366935483870968 |
|  | 54 | CAG | 177 | 0.356854838709677 |
|  | 55 | CAC | 226 | 0.455645161290323 |
|  | 56 | CAU | 229 | 0.461693548387097 |
|  | 57 | CCA | 203 | 0.409274193548387 |
|  | 58 | CCG | 262 | 0.528225806451613 |
|  | 59 | CCC | 215 | 0.433467741935484 |
|  | 60 | CCU | 231 | 0.465725806451613 |
|  | 61 | CGA | 210 | 0.423387096774194 |
|  | 62 | CGG | 270 | 0.544354838709677 |
|  | 63 | CGC | 264 | 0.532258064516129 |
|  | 64 | CGU | 217 | 0.437500000000000 |
|  | 65 | CUA | 253 | 0.510080645161290 |
|  | 66 | CUG | 200 | 0.403225806451613 |
|  | 67 | CUC | 261 | 0.526209677419355 |
|  | 68 | CUU | 157 | 0.316532258064516 |
|  | 69 | UAA | 268 | 0.540322580645161 |
|  | 70 | UAG | 219 | 0.441532258064516 |
|  | 71 | UAC | 132 | 0.266129032258065 |
|  | 72 | UAU | 258 | 0.520161290322581 |
|  | 73 | UCA | 214 | 0.431451612903226 |
|  | 74 | UCG | 211 | 0.425403225806452 |
|  | 75 | UCC | 266 | 0.536290322580645 |
|  | 76 | UCU | 156 | 0.314516129032258 |
|  | 77 | UGA | 154 | 0.310483870967742 |
|  | 78 | UGG | 226 | 0.455645161290323 |
|  | 79 | UGC | 224 | 0.451612903225806 |
|  | 80 | UGU | 221 | 0.445564516129032 |
|  | 81 | UUA | 269 | 0.542338709677419 |
|  | 82 | UUG | 196 | 0.395161290322581 |
|  | 83 | UUC | 214 | 0.431451612903226 |
|  | 84 | UUU | 254 | 0.512096774193548 |
| D | 85 | A... | 267 | 0.538306451612903 |
|  | 86 | A..+ | 139 | 0.280241935483871 |
|  | 87 | A.+. | 180 | 0.362903225806452 |
|  | 88 | A+.. | 55 | 0.110887096774194 |
|  | 89 | A.++ | 50 | 0.100806451612903 |
|  | 90 | A+.+ | 176 | 0.354838709677419 |
|  | 91 | A++. | 111 | 0.223790322580645 |
|  | 92 | A+++ | 128 | 0.258064516129032 |
|  | 93 | C... | 226 | 0.455645161290323 |
|  | 94 | C..+ | 184 | 0.370967741935484 |
|  | 95 | C.+. | 203 | 0.409274193548387 |
|  | 96 | C+.. | 134 | 0.270161290322581 |
|  | 97 | C.++ | 136 | 0.274193548387097 |
|  | 98 | C+.+ | 188 | 0.379032258064516 |
|  | 99 | C++. | 118 | 0.237903225806452 |
|  | 100 | C+++ | 116 | 0.233870967741935 |
|  | 101 | G... | 207 | 0.417338709677419 |
|  | 102 | G..+ | 189 | 0.381048387096774 |
|  | 103 | G.+. | 158 | 0.318548387096774 |
|  | 104 | G+.. | 122 | 0.245967741935484 |
|  | 105 | G.++ | 106 | 0.213709677419355 |
|  | 106 | G+.+ | 205 | 0.413306451612903 |
|  | 107 | G++. | 171 | 0.344758064516129 |
|  | 108 | G+++ | 182 | 0.366935483870968 |
|  | 109 | U... | 233 | 0.469758064516129 |
|  | 110 | U..+ | 156 | 0.314516129032258 |
|  | 111 | U.+. | 162 | 0.326612903225806 |
|  | 112 | U+.. | 69 | 0.139112903225806 |
|  | 113 | U.++ | 80 | 0.161290322580645 |
|  | 114 | U+.+ | 146 | 0.294354838709677 |
|  | 115 | U++. | 84 | 0.169354838709677 |
|  | 116 | U+++ | 130 | 0.262096774193548 |
| E | 117 | Bulge loop | 71 | 0.143145161290323 |
|  | 118 | Helix | 323 | 0.651209677419355 |
|  | 119 | Inter loop | 257 | 0.518145161290323 |
|  | 120 | Stack | 364 | 0.733870967741936 |
|  | 121 | Length | 445 | 0.897177419354839 |
| F | 122 | MFE | 375 | 0.756048387096774 |
|  | 123 | AMFE | 307 | 0.618951612903226 |
|  | 124 | MFEI | 301 | 0.606854838709677 |
| G | 125 | G+C | 357 | 0.719758064516129 |
|  | 126 | (G+C)/(A+U) | 358 | 0.721774193548387 |
|  | 127 | A/C | 323 | 0.651209677419355 |
|  | 128 | G/U | 328 | 0.661290322580645 |
| H | 129 | One entroy | 190 | 0.383064516129032 |
|  | 130 | Two entroy | 194 | 0.391129032258065 |
|  | 131 | Threeentroy | 349 | 0.703629032258065 |
|  | 132 | Sec_str_entr | 230 | 0.463709677419355 |

FIN-Feature identification number

FIR- Feature identification ratio;FIN/496
