## Supplementary Table 2 for "Reconstructing species evolutionary trees using the numerical features of microRNA"

Table S2 State of species pairs being identified

| one Species | | Another species | | SPIN | | SPIR |
| --- | --- | --- | --- | --- | --- | --- |
| 1 | 2 | | 77 | | 0.583333333333333 | |
| 1 | 3 | | 116 | | 0.878787878787879 | |
| 1 | 4 | | 69 | | 0.522727272727273 | |
| 1 | 5 | | 85 | | 0.643939393939394 | |
| 1 | 6 | | 75 | | 0.568181818181818 | |
| 1 | 7 | | 111 | | 0.840909090909091 | |
| 1 | 8 | | 113 | | 0.856060606060606 | |
| 1 | 9 | | 111 | | 0.840909090909091 | |
| 1 | 10 | | 114 | | 0.863636363636364 | |
| 1 | 11 | | 117 | | 0.886363636363636 | |
| 1 | 12 | | 110 | | 0.833333333333333 | |
| 1 | 13 | | 107 | | 0.810606060606061 | |
| 1 | 14 | | 114 | | 0.863636363636364 | |
| 1 | 15 | | 99 | | 0.750000000000000 | |
| 1 | 16 | | 111 | | 0.840909090909091 | |
| 1 | 17 | | 72 | | 0.545454545454545 | |
| 1 | 18 | | 99 | | 0.750000000000000 | |
| 1 | 19 | | 48 | | 0.363636363636364 | |
| 1 | 20 | | 48 | | 0.363636363636364 | |
| 1 | 21 | | 72 | | 0.545454545454545 | |
| 1 | 22 | | 74 | | 0.560606060606061 | |
| 1 | 23 | | 125 | | 0.946969696969697 | |
| 1 | 24 | | 108 | | 0.818181818181818 | |
| 1 | 25 | | 70 | | 0.530303030303030 | |
| 1 | 26 | | 119 | | 0.901515151515152 | |
| 1 | 27 | | 90 | | 0.681818181818182 | |
| 1 | 28 | | 98 | | 0.742424242424242 | |
| 1 | 29 | | 119 | | 0.901515151515152 | |
| 1 | 30 | | 86 | | 0.651515151515152 | |
| 1 | 31 | | 110 | | 0.833333333333333 | |
| 1 | 32 | | 106 | | 0.803030303030303 | |
| 2 | 3 | | 41 | | 0.310606060606061 | |
| 2 | 4 | | 17 | | 0.128787878787879 | |
| 2 | 5 | | 4 | | 0.0303030303030303 | |
| 2 | 6 | | 0 | | 0 | |
| 2 | 7 | | 18 | | 0.136363636363636 | |
| 2 | 8 | | 47 | | 0.356060606060606 | |
| 2 | 9 | | 19 | | 0.143939393939394 | |
| 2 | 10 | | 25 | | 0.189393939393939 | |
| 2 | 11 | | 51 | | 0.386363636363636 | |
| 2 | 12 | | 23 | | 0.174242424242424 | |
| 2 | 13 | | 26 | | 0.196969696969697 | |
| 2 | 14 | | 47 | | 0.356060606060606 | |
| 2 | 15 | | 14 | | 0.106060606060606 | |
| 2 | 16 | | 7 | | 0.0530303030303030 | |
| 2 | 17 | | 0 | | 0 | |
| 2 | 18 | | 51 | | 0.386363636363636 | |
| 2 | 19 | | 34 | | 0.257575757575758 | |
| 2 | 20 | | 48 | | 0.363636363636364 | |
| 2 | 21 | | 16 | | 0.121212121212121 | |
| 2 | 22 | | 74 | | 0.560606060606061 | |
| 2 | 23 | | 59 | | 0.446969696969697 | |
| 2 | 24 | | 94 | | 0.712121212121212 | |
| 2 | 25 | | 74 | | 0.560606060606061 | |
| 2 | 26 | | 98 | | 0.742424242424242 | |
| 2 | 27 | | 45 | | 0.340909090909091 | |
| 2 | 28 | | 37 | | 0.280303030303030 | |
| 2 | 29 | | 72 | | 0.545454545454545 | |
| 2 | 30 | | 41 | | 0.310606060606061 | |
| 2 | 31 | | 77 | | 0.583333333333333 | |
| 2 | 32 | | 73 | | 0.553030303030303 | |
| 3 | 4 | | 63 | | 0.477272727272727 | |
| 3 | 5 | | 35 | | 0.265151515151515 | |
| 3 | 6 | | 19 | | 0.143939393939394 | |
| 3 | 7 | | 12 | | 0.0909090909090909 | |
| 3 | 8 | | 31 | | 0.234848484848485 | |
| 3 | 9 | | 23 | | 0.174242424242424 | |
| 3 | 10 | | 12 | | 0.0909090909090909 | |
| 3 | 11 | | 30 | | 0.227272727272727 | |
| 3 | 12 | | 20 | | 0.151515151515152 | |
| 3 | 13 | | 14 | | 0.106060606060606 | |
| 3 | 14 | | 19 | | 0.143939393939394 | |
| 3 | 15 | | 23 | | 0.174242424242424 | |
| 3 | 16 | | 65 | | 0.492424242424242 | |
| 3 | 17 | | 28 | | 0.212121212121212 | |
| 3 | 18 | | 80 | | 0.606060606060606 | |
| 3 | 19 | | 62 | | 0.469696969696970 | |
| 3 | 20 | | 68 | | 0.515151515151515 | |
| 3 | 21 | | 33 | | 0.250000000000000 | |
| 3 | 22 | | 83 | | 0.628787878787879 | |
| 3 | 23 | | 101 | | 0.765151515151515 | |
| 3 | 24 | | 112 | | 0.848484848484849 | |
| 3 | 25 | | 98 | | 0.742424242424242 | |
| 3 | 26 | | 118 | | 0.893939393939394 | |
| 3 | 27 | | 83 | | 0.628787878787879 | |
| 3 | 28 | | 68 | | 0.515151515151515 | |
| 3 | 29 | | 102 | | 0.772727272727273 | |
| 3 | 30 | | 38 | | 0.287878787878788 | |
| 3 | 31 | | 64 | | 0.484848484848485 | |
| 3 | 32 | | 54 | | 0.409090909090909 | |
| 4 | 5 | | 6 | | 0.0454545454545455 | |
| 4 | 6 | | 6 | | 0.0454545454545455 | |
| 4 | 7 | | 24 | | 0.181818181818182 | |
| 4 | 8 | | 54 | | 0.409090909090909 | |
| 4 | 9 | | 38 | | 0.287878787878788 | |
| 4 | 10 | | 36 | | 0.272727272727273 | |
| 4 | 11 | | 59 | | 0.446969696969697 | |
| 4 | 12 | | 33 | | 0.250000000000000 | |
| 4 | 13 | | 30 | | 0.227272727272727 | |
| 4 | 14 | | 60 | | 0.454545454545455 | |
| 4 | 15 | | 18 | | 0.136363636363636 | |
| 4 | 16 | | 80 | | 0.606060606060606 | |
| 4 | 17 | | 11 | | 0.0833333333333333 | |
| 4 | 18 | | 99 | | 0.750000000000000 | |
| 4 | 19 | | 65 | | 0.492424242424242 | |
| 4 | 20 | | 60 | | 0.454545454545455 | |
| 4 | 21 | | 29 | | 0.219696969696970 | |
| 4 | 22 | | 94 | | 0.712121212121212 | |
| 4 | 23 | | 102 | | 0.772727272727273 | |
| 4 | 24 | | 122 | | 0.924242424242424 | |
| 4 | 25 | | 86 | | 0.651515151515152 | |
| 4 | 26 | | 122 | | 0.924242424242424 | |
| 4 | 27 | | 85 | | 0.643939393939394 | |
| 4 | 28 | | 69 | | 0.522727272727273 | |
| 4 | 29 | | 105 | | 0.795454545454545 | |
| 4 | 30 | | 49 | | 0.371212121212121 | |
| 4 | 31 | | 87 | | 0.659090909090909 | |
| 4 | 32 | | 74 | | 0.560606060606061 | |
| 5 | 6 | | 0 | | 0 | |
| 5 | 7 | | 6 | | 0.0454545454545455 | |
| 5 | 8 | | 23 | | 0.174242424242424 | |
| 5 | 9 | | 7 | | 0.0530303030303030 | |
| 5 | 10 | | 7 | | 0.0530303030303030 | |
| 5 | 11 | | 41 | | 0.310606060606061 | |
| 5 | 12 | | 12 | | 0.0909090909090909 | |
| 5 | 13 | | 15 | | 0.113636363636364 | |
| 5 | 14 | | 20 | | 0.151515151515152 | |
| 5 | 15 | | 1 | | 0.00757575757575758 | |
| 5 | 16 | | 44 | | 0.333333333333333 | |
| 5 | 17 | | 2 | | 0.0151515151515152 | |
| 5 | 18 | | 70 | | 0.530303030303030 | |
| 5 | 19 | | 50 | | 0.378787878787879 | |
| 5 | 20 | | 56 | | 0.424242424242424 | |
| 5 | 21 | | 21 | | 0.159090909090909 | |
| 5 | 22 | | 86 | | 0.651515151515152 | |
| 5 | 23 | | 90 | | 0.681818181818182 | |
| 5 | 24 | | 115 | | 0.871212121212121 | |
| 5 | 25 | | 80 | | 0.606060606060606 | |
| 5 | 26 | | 119 | | 0.901515151515152 | |
| 5 | 27 | | 66 | | 0.500000000000000 | |
| 5 | 28 | | 53 | | 0.401515151515152 | |
| 5 | 29 | | 84 | | 0.636363636363636 | |
| 5 | 30 | | 30 | | 0.227272727272727 | |
| 5 | 31 | | 70 | | 0.530303030303030 | |
| 5 | 32 | | 54 | | 0.409090909090909 | |
| 6 | 7 | | 3 | | 0.0227272727272727 | |
| 6 | 8 | | 19 | | 0.143939393939394 | |
| 6 | 9 | | 3 | | 0.0227272727272727 | |
| 6 | 10 | | 4 | | 0.0303030303030303 | |
| 6 | 11 | | 36 | | 0.272727272727273 | |
| 6 | 12 | | 6 | | 0.0454545454545455 | |
| 6 | 13 | | 9 | | 0.0681818181818182 | |
| 6 | 14 | | 20 | | 0.151515151515152 | |
| 6 | 15 | | 2 | | 0.0151515151515152 | |
| 6 | 16 | | 12 | | 0.0909090909090909 | |
| 6 | 17 | | 0 | | 0 | |
| 6 | 18 | | 58 | | 0.439393939393939 | |
| 6 | 19 | | 38 | | 0.287878787878788 | |
| 6 | 20 | | 48 | | 0.363636363636364 | |
| 6 | 21 | | 18 | | 0.136363636363636 | |
| 6 | 22 | | 74 | | 0.560606060606061 | |
| 6 | 23 | | 53 | | 0.401515151515152 | |
| 6 | 24 | | 91 | | 0.689393939393939 | |
| 6 | 25 | | 65 | | 0.492424242424242 | |
| 6 | 26 | | 91 | | 0.689393939393939 | |
| 6 | 27 | | 41 | | 0.310606060606061 | |
| 6 | 28 | | 35 | | 0.265151515151515 | |
| 6 | 29 | | 53 | | 0.401515151515152 | |
| 6 | 30 | | 24 | | 0.181818181818182 | |
| 6 | 31 | | 61 | | 0.462121212121212 | |
| 6 | 32 | | 49 | | 0.371212121212121 | |
| 7 | 8 | | 2 | | 0.0151515151515152 | |
| 7 | 9 | | 0 | | 0 | |
| 7 | 10 | | 0 | | 0 | |
| 7 | 11 | | 30 | | 0.227272727272727 | |
| 7 | 12 | | 4 | | 0.0303030303030303 | |
| 7 | 13 | | 1 | | 0.00757575757575758 | |
| 7 | 14 | | 12 | | 0.0909090909090909 | |
| 7 | 15 | | 5 | | 0.0378787878787879 | |
| 7 | 16 | | 49 | | 0.371212121212121 | |
| 7 | 17 | | 8 | | 0.0606060606060606 | |
| 7 | 18 | | 75 | | 0.568181818181818 | |
| 7 | 19 | | 56 | | 0.424242424242424 | |
| 7 | 20 | | 63 | | 0.477272727272727 | |
| 7 | 21 | | 24 | | 0.181818181818182 | |
| 7 | 22 | | 84 | | 0.636363636363636 | |
| 7 | 23 | | 93 | | 0.704545454545455 | |
| 7 | 24 | | 110 | | 0.833333333333333 | |
| 7 | 25 | | 90 | | 0.681818181818182 | |
| 7 | 26 | | 117 | | 0.886363636363636 | |
| 7 | 27 | | 71 | | 0.537878787878788 | |
| 7 | 28 | | 49 | | 0.371212121212121 | |
| 7 | 29 | | 92 | | 0.696969696969697 | |
| 7 | 30 | | 29 | | 0.219696969696970 | |
| 7 | 31 | | 69 | | 0.522727272727273 | |
| 7 | 32 | | 57 | | 0.431818181818182 | |
| 8 | 9 | | 3 | | 0.0227272727272727 | |
| 8 | 10 | | 4 | | 0.0303030303030303 | |
| 8 | 11 | | 58 | | 0.439393939393939 | |
| 8 | 12 | | 28 | | 0.212121212121212 | |
| 8 | 13 | | 17 | | 0.128787878787879 | |
| 8 | 14 | | 20 | | 0.151515151515152 | |
| 8 | 15 | | 17 | | 0.128787878787879 | |
| 8 | 16 | | 78 | | 0.590909090909091 | |
| 8 | 17 | | 32 | | 0.242424242424242 | |
| 8 | 18 | | 91 | | 0.689393939393939 | |
| 8 | 19 | | 73 | | 0.553030303030303 | |
| 8 | 20 | | 77 | | 0.583333333333333 | |
| 8 | 21 | | 49 | | 0.371212121212121 | |
| 8 | 22 | | 86 | | 0.651515151515152 | |
| 8 | 23 | | 119 | | 0.901515151515152 | |
| 8 | 24 | | 123 | | 0.931818181818182 | |
| 8 | 25 | | 93 | | 0.704545454545455 | |
| 8 | 26 | | 125 | | 0.946969696969697 | |
| 8 | 27 | | 95 | | 0.719696969696970 | |
| 8 | 28 | | 65 | | 0.492424242424242 | |
| 8 | 29 | | 107 | | 0.810606060606061 | |
| 8 | 30 | | 35 | | 0.265151515151515 | |
| 8 | 31 | | 80 | | 0.606060606060606 | |
| 8 | 32 | | 57 | | 0.431818181818182 | |
| 9 | 10 | | 0 | | 0 | |
| 9 | 11 | | 51 | | 0.386363636363636 | |
| 9 | 12 | | 16 | | 0.121212121212121 | |
| 9 | 13 | | 9 | | 0.0681818181818182 | |
| 9 | 14 | | 26 | | 0.196969696969697 | |
| 9 | 15 | | 10 | | 0.0757575757575758 | |
| 9 | 16 | | 49 | | 0.371212121212121 | |
| 9 | 17 | | 12 | | 0.0909090909090909 | |
| 9 | 18 | | 71 | | 0.537878787878788 | |
| 9 | 19 | | 54 | | 0.409090909090909 | |
| 9 | 20 | | 60 | | 0.454545454545455 | |
| 9 | 21 | | 27 | | 0.204545454545455 | |
| 9 | 22 | | 81 | | 0.613636363636364 | |
| 9 | 23 | | 95 | | 0.719696969696970 | |
| 9 | 24 | | 111 | | 0.840909090909091 | |
| 9 | 25 | | 89 | | 0.674242424242424 | |
| 9 | 26 | | 117 | | 0.886363636363636 | |
| 9 | 27 | | 70 | | 0.530303030303030 | |
| 9 | 28 | | 50 | | 0.378787878787879 | |
| 9 | 29 | | 92 | | 0.696969696969697 | |
| 9 | 30 | | 35 | | 0.265151515151515 | |
| 9 | 31 | | 72 | | 0.545454545454545 | |
| 9 | 32 | | 64 | | 0.484848484848485 | |
| 10 | 11 | | 37 | | 0.280303030303030 | |
| 10 | 12 | | 11 | | 0.0833333333333333 | |
| 10 | 13 | | 2 | | 0.0151515151515152 | |
| 10 | 14 | | 15 | | 0.113636363636364 | |
| 10 | 15 | | 10 | | 0.0757575757575758 | |
| 10 | 16 | | 54 | | 0.409090909090909 | |
| 10 | 17 | | 11 | | 0.0833333333333333 | |
| 10 | 18 | | 77 | | 0.583333333333333 | |
| 10 | 19 | | 62 | | 0.469696969696970 | |
| 10 | 20 | | 63 | | 0.477272727272727 | |
| 10 | 21 | | 27 | | 0.204545454545455 | |
| 10 | 22 | | 82 | | 0.621212121212121 | |
| 10 | 23 | | 95 | | 0.719696969696970 | |
| 10 | 24 | | 114 | | 0.863636363636364 | |
| 10 | 25 | | 88 | | 0.666666666666667 | |
| 10 | 26 | | 116 | | 0.878787878787879 | |
| 10 | 27 | | 72 | | 0.545454545454545 | |
| 10 | 28 | | 51 | | 0.386363636363636 | |
| 10 | 29 | | 95 | | 0.719696969696970 | |
| 10 | 30 | | 35 | | 0.265151515151515 | |
| 10 | 31 | | 67 | | 0.507575757575758 | |
| 10 | 32 | | 60 | | 0.454545454545455 | |
| 11 | 12 | | 39 | | 0.295454545454545 | |
| 11 | 13 | | 14 | | 0.106060606060606 | |
| 11 | 14 | | 41 | | 0.310606060606061 | |
| 11 | 15 | | 31 | | 0.234848484848485 | |
| 11 | 16 | | 68 | | 0.515151515151515 | |
| 11 | 17 | | 38 | | 0.287878787878788 | |
| 11 | 18 | | 70 | | 0.530303030303030 | |
| 11 | 19 | | 50 | | 0.378787878787879 | |
| 11 | 20 | | 71 | | 0.537878787878788 | |
| 11 | 21 | | 34 | | 0.257575757575758 | |
| 11 | 22 | | 98 | | 0.742424242424242 | |
| 11 | 23 | | 54 | | 0.409090909090909 | |
| 11 | 24 | | 95 | | 0.719696969696970 | |
| 11 | 25 | | 85 | | 0.643939393939394 | |
| 11 | 26 | | 101 | | 0.765151515151515 | |
| 11 | 27 | | 63 | | 0.477272727272727 | |
| 11 | 28 | | 44 | | 0.333333333333333 | |
| 11 | 29 | | 82 | | 0.621212121212121 | |
| 11 | 30 | | 30 | | 0.227272727272727 | |
| 11 | 31 | | 33 | | 0.250000000000000 | |
| 11 | 32 | | 64 | | 0.484848484848485 | |
| 12 | 13 | | 4 | | 0.0303030303030303 | |
| 12 | 14 | | 32 | | 0.242424242424242 | |
| 12 | 15 | | 11 | | 0.0833333333333333 | |
| 12 | 16 | | 69 | | 0.522727272727273 | |
| 12 | 17 | | 11 | | 0.0833333333333333 | |
| 12 | 18 | | 79 | | 0.598484848484849 | |
| 12 | 19 | | 64 | | 0.484848484848485 | |
| 12 | 20 | | 62 | | 0.469696969696970 | |
| 12 | 21 | | 26 | | 0.196969696969697 | |
| 12 | 22 | | 86 | | 0.651515151515152 | |
| 12 | 23 | | 107 | | 0.810606060606061 | |
| 12 | 24 | | 120 | | 0.909090909090909 | |
| 12 | 25 | | 91 | | 0.689393939393939 | |
| 12 | 26 | | 123 | | 0.931818181818182 | |
| 12 | 27 | | 79 | | 0.598484848484849 | |
| 12 | 28 | | 60 | | 0.454545454545455 | |
| 12 | 29 | | 111 | | 0.840909090909091 | |
| 12 | 30 | | 40 | | 0.303030303030303 | |
| 12 | 31 | | 79 | | 0.598484848484849 | |
| 12 | 32 | | 59 | | 0.446969696969697 | |
| 13 | 14 | | 16 | | 0.121212121212121 | |
| 13 | 15 | | 5 | | 0.0378787878787879 | |
| 13 | 16 | | 45 | | 0.340909090909091 | |
| 13 | 17 | | 16 | | 0.121212121212121 | |
| 13 | 18 | | 68 | | 0.515151515151515 | |
| 13 | 19 | | 47 | | 0.356060606060606 | |
| 13 | 20 | | 60 | | 0.454545454545455 | |
| 13 | 21 | | 24 | | 0.181818181818182 | |
| 13 | 22 | | 85 | | 0.643939393939394 | |
| 13 | 23 | | 73 | | 0.553030303030303 | |
| 13 | 24 | | 105 | | 0.795454545454545 | |
| 13 | 25 | | 87 | | 0.659090909090909 | |
| 13 | 26 | | 118 | | 0.893939393939394 | |
| 13 | 27 | | 64 | | 0.484848484848485 | |
| 13 | 28 | | 47 | | 0.356060606060606 | |
| 13 | 29 | | 95 | | 0.719696969696970 | |
| 13 | 30 | | 32 | | 0.242424242424242 | |
| 13 | 31 | | 56 | | 0.424242424242424 | |
| 13 | 32 | | 55 | | 0.416666666666667 | |
| 14 | 15 | | 11 | | 0.0833333333333333 | |
| 14 | 16 | | 93 | | 0.704545454545455 | |
| 14 | 17 | | 29 | | 0.219696969696970 | |
| 14 | 18 | | 89 | | 0.674242424242424 | |
| 14 | 19 | | 78 | | 0.590909090909091 | |
| 14 | 20 | | 81 | | 0.613636363636364 | |
| 14 | 21 | | 38 | | 0.287878787878788 | |
| 14 | 22 | | 93 | | 0.704545454545455 | |
| 14 | 23 | | 111 | | 0.840909090909091 | |
| 14 | 24 | | 120 | | 0.909090909090909 | |
| 14 | 25 | | 100 | | 0.757575757575758 | |
| 14 | 26 | | 121 | | 0.916666666666667 | |
| 14 | 27 | | 99 | | 0.750000000000000 | |
| 14 | 28 | | 82 | | 0.621212121212121 | |
| 14 | 29 | | 116 | | 0.878787878787879 | |
| 14 | 30 | | 50 | | 0.378787878787879 | |
| 14 | 31 | | 79 | | 0.598484848484849 | |
| 14 | 32 | | 38 | | 0.287878787878788 | |
| 15 | 16 | | 47 | | 0.356060606060606 | |
| 15 | 17 | | 3 | | 0.0227272727272727 | |
| 15 | 18 | | 78 | | 0.590909090909091 | |
| 15 | 19 | | 56 | | 0.424242424242424 | |
| 15 | 20 | | 67 | | 0.507575757575758 | |
| 15 | 21 | | 18 | | 0.136363636363636 | |
| 15 | 22 | | 90 | | 0.681818181818182 | |
| 15 | 23 | | 90 | | 0.681818181818182 | |
| 15 | 24 | | 112 | | 0.848484848484849 | |
| 15 | 25 | | 92 | | 0.696969696969697 | |
| 15 | 26 | | 119 | | 0.901515151515152 | |
| 15 | 27 | | 74 | | 0.560606060606061 | |
| 15 | 28 | | 55 | | 0.416666666666667 | |
| 15 | 29 | | 94 | | 0.712121212121212 | |
| 15 | 30 | | 39 | | 0.295454545454545 | |
| 15 | 31 | | 69 | | 0.522727272727273 | |
| 15 | 32 | | 47 | | 0.356060606060606 | |
| 16 | 17 | | 7 | | 0.0530303030303030 | |
| 16 | 18 | | 65 | | 0.492424242424242 | |
| 16 | 19 | | 44 | | 0.333333333333333 | |
| 16 | 20 | | 43 | | 0.325757575757576 | |
| 16 | 21 | | 18 | | 0.136363636363636 | |
| 16 | 22 | | 82 | | 0.621212121212121 | |
| 16 | 23 | | 80 | | 0.606060606060606 | |
| 16 | 24 | | 86 | | 0.651515151515152 | |
| 16 | 25 | | 80 | | 0.606060606060606 | |
| 16 | 26 | | 107 | | 0.810606060606061 | |
| 16 | 27 | | 49 | | 0.371212121212121 | |
| 16 | 28 | | 48 | | 0.363636363636364 | |
| 16 | 29 | | 92 | | 0.696969696969697 | |
| 16 | 30 | | 54 | | 0.409090909090909 | |
| 16 | 31 | | 81 | | 0.613636363636364 | |
| 16 | 32 | | 94 | | 0.712121212121212 | |
| 17 | 18 | | 54 | | 0.409090909090909 | |
| 17 | 19 | | 44 | | 0.333333333333333 | |
| 17 | 20 | | 43 | | 0.325757575757576 | |
| 17 | 21 | | 16 | | 0.121212121212121 | |
| 17 | 22 | | 73 | | 0.553030303030303 | |
| 17 | 23 | | 52 | | 0.393939393939394 | |
| 17 | 24 | | 84 | | 0.636363636363636 | |
| 17 | 25 | | 70 | | 0.530303030303030 | |
| 17 | 26 | | 93 | | 0.704545454545455 | |
| 17 | 27 | | 42 | | 0.318181818181818 | |
| 17 | 28 | | 35 | | 0.265151515151515 | |
| 17 | 29 | | 60 | | 0.454545454545455 | |
| 17 | 30 | | 33 | | 0.250000000000000 | |
| 17 | 31 | | 66 | | 0.500000000000000 | |
| 17 | 32 | | 55 | | 0.416666666666667 | |
| 18 | 19 | | 12 | | 0.0909090909090909 | |
| 18 | 20 | | 19 | | 0.143939393939394 | |
| 18 | 21 | | 34 | | 0.257575757575758 | |
| 18 | 22 | | 76 | | 0.575757575757576 | |
| 18 | 23 | | 88 | | 0.666666666666667 | |
| 18 | 24 | | 80 | | 0.606060606060606 | |
| 18 | 25 | | 59 | | 0.446969696969697 | |
| 18 | 26 | | 96 | | 0.727272727272727 | |
| 18 | 27 | | 58 | | 0.439393939393939 | |
| 18 | 28 | | 65 | | 0.492424242424242 | |
| 18 | 29 | | 67 | | 0.507575757575758 | |
| 18 | 30 | | 50 | | 0.378787878787879 | |
| 18 | 31 | | 83 | | 0.628787878787879 | |
| 18 | 32 | | 80 | | 0.606060606060606 | |
| 19 | 20 | | 6 | | 0.0454545454545455 | |
| 19 | 21 | | 22 | | 0.166666666666667 | |
| 19 | 22 | | 53 | | 0.401515151515152 | |
| 19 | 23 | | 58 | | 0.439393939393939 | |
| 19 | 24 | | 53 | | 0.401515151515152 | |
| 19 | 25 | | 45 | | 0.340909090909091 | |
| 19 | 26 | | 79 | | 0.598484848484849 | |
| 19 | 27 | | 35 | | 0.265151515151515 | |
| 19 | 28 | | 44 | | 0.333333333333333 | |
| 19 | 29 | | 41 | | 0.310606060606061 | |
| 19 | 30 | | 39 | | 0.295454545454545 | |
| 19 | 31 | | 74 | | 0.560606060606061 | |
| 19 | 32 | | 79 | | 0.598484848484849 | |
| 20 | 21 | | 20 | | 0.151515151515152 | |
| 20 | 22 | | 49 | | 0.371212121212121 | |
| 20 | 23 | | 65 | | 0.492424242424242 | |
| 20 | 24 | | 54 | | 0.409090909090909 | |
| 20 | 25 | | 36 | | 0.272727272727273 | |
| 20 | 26 | | 86 | | 0.651515151515152 | |
| 20 | 27 | | 30 | | 0.227272727272727 | |
| 20 | 28 | | 38 | | 0.287878787878788 | |
| 20 | 29 | | 48 | | 0.363636363636364 | |
| 20 | 30 | | 43 | | 0.325757575757576 | |
| 20 | 31 | | 75 | | 0.568181818181818 | |
| 20 | 32 | | 81 | | 0.613636363636364 | |
| 21 | 22 | | 68 | | 0.515151515151515 | |
| 21 | 23 | | 25 | | 0.189393939393939 | |
| 21 | 24 | | 74 | | 0.560606060606061 | |
| 21 | 25 | | 64 | | 0.484848484848485 | |
| 21 | 26 | | 90 | | 0.681818181818182 | |
| 21 | 27 | | 38 | | 0.287878787878788 | |
| 21 | 28 | | 22 | | 0.166666666666667 | |
| 21 | 29 | | 51 | | 0.386363636363636 | |
| 21 | 30 | | 22 | | 0.166666666666667 | |
| 21 | 31 | | 56 | | 0.424242424242424 | |
| 21 | 32 | | 53 | | 0.401515151515152 | |
| 22 | 23 | | 95 | | 0.719696969696970 | |
| 22 | 24 | | 51 | | 0.386363636363636 | |
| 22 | 25 | | 47 | | 0.356060606060606 | |
| 22 | 26 | | 49 | | 0.371212121212121 | |
| 22 | 27 | | 60 | | 0.454545454545455 | |
| 22 | 28 | | 72 | | 0.545454545454545 | |
| 22 | 29 | | 73 | | 0.553030303030303 | |
| 22 | 30 | | 71 | | 0.537878787878788 | |
| 22 | 31 | | 96 | | 0.727272727272727 | |
| 22 | 32 | | 103 | | 0.780303030303030 | |
| 23 | 24 | | 66 | | 0.500000000000000 | |
| 23 | 25 | | 73 | | 0.553030303030303 | |
| 23 | 26 | | 85 | | 0.643939393939394 | |
| 23 | 27 | | 43 | | 0.325757575757576 | |
| 23 | 28 | | 19 | | 0.143939393939394 | |
| 23 | 29 | | 58 | | 0.439393939393939 | |
| 23 | 30 | | 28 | | 0.212121212121212 | |
| 23 | 31 | | 42 | | 0.318181818181818 | |
| 23 | 32 | | 103 | | 0.780303030303030 | |
| 24 | 25 | | 21 | | 0.159090909090909 | |
| 24 | 26 | | 43 | | 0.325757575757576 | |
| 24 | 27 | | 45 | | 0.340909090909091 | |
| 24 | 28 | | 58 | | 0.439393939393939 | |
| 24 | 29 | | 62 | | 0.469696969696970 | |
| 24 | 30 | | 48 | | 0.363636363636364 | |
| 24 | 31 | | 89 | | 0.674242424242424 | |
| 24 | 32 | | 116 | | 0.878787878787879 | |
| 25 | 26 | | 45 | | 0.340909090909091 | |
| 25 | 27 | | 44 | | 0.333333333333333 | |
| 25 | 28 | | 47 | | 0.356060606060606 | |
| 25 | 29 | | 31 | | 0.234848484848485 | |
| 25 | 30 | | 47 | | 0.356060606060606 | |
| 25 | 31 | | 76 | | 0.575757575757576 | |
| 25 | 32 | | 96 | | 0.727272727272727 | |
| 26 | 27 | | 70 | | 0.530303030303030 | |
| 26 | 28 | | 66 | | 0.500000000000000 | |
| 26 | 29 | | 78 | | 0.590909090909091 | |
| 26 | 30 | | 68 | | 0.515151515151515 | |
| 26 | 31 | | 92 | | 0.696969696969697 | |
| 26 | 32 | | 122 | | 0.924242424242424 | |
| 27 | 28 | | 4 | | 0.0303030303030303 | |
| 27 | 29 | | 38 | | 0.287878787878788 | |
| 27 | 30 | | 25 | | 0.189393939393939 | |
| 27 | 31 | | 66 | | 0.500000000000000 | |
| 27 | 32 | | 97 | | 0.734848484848485 | |
| 28 | 29 | | 37 | | 0.280303030303030 | |
| 28 | 30 | | 9 | | 0.0681818181818182 | |
| 28 | 31 | | 47 | | 0.356060606060606 | |
| 28 | 32 | | 89 | | 0.674242424242424 | |
| 29 | 30 | | 14 | | 0.106060606060606 | |
| 29 | 31 | | 62 | | 0.469696969696970 | |
| 29 | 32 | | 96 | | 0.727272727272727 | |
| 30 | 31 | | 9 | | 0.0681818181818182 | |
| 30 | 32 | | 68 | | 0.515151515151515 | |
| 31 | 32 | | 59 | | 0.446969696969697 | |

The numbers of the two first columns denotes: 1-Ciona intestinalis, 2-Xenopus tropicalis, 3-Gallus gallus, 4-Canis familiaris, 5-Equus caballus, 6-Monodelphis domestica, 7-Macaca mulatta, 8-Homo sapiens, 9-Pan troglodytes, 10-Pongo pygmaeus, 11-Ornithorhynchus anatinus, 12-Mus musculus, 13-Rattus norvegicus, 14-Bos taurus, 15-Sus scrofa, 16-Danio rerio, 17-Fugu rubripes, 18-Bombyx mori, 19-Drosophila pseudoobscura, 20-Caenorhabditis elegans, 21-Capitella teleta, 22-Schmidtea mediterranea, 23-Physcomitrella patens, 24-Arabidopsis thaliana, 25-Glycine max, 26-Medicago truncatula, 27-Populus trichocarpa, 28-Vitis vinifera, 29-Oryza sativa, 30-Sorghum bicolor, 31-Zea mays, 32-Virus
